# Characterization of an EG.5.1 clinical isolate *in vitro* and *in vivo*

**DOI:** 10.1101/2023.08.31.555819

**Authors:** Ryuta Uraki, Maki Kiso, Kiyoko Iwatsuki-Horimoto, Seiya Yamayoshi, Mutsumi Ito, Shiho Chiba, Yuko Sakai-Tagawa, Masaki Imai, Yukie Kashima, Michiko Koga, Noriko Fuwa, Nobumasa Okumura, Masayuki Hojo, Noriko Iwamoto, Hideaki Kato, Hideaki Nakajima, Norio Ohmagari, Hiroshi Yotsuyanagi, Yutaka Suzuki, Yoshihiro Kawaoka

## Abstract

EG.5.1 is a subvariant of the SARS-CoV-2 Omicron XBB variant that is rapidly increasing in prevalence worldwide. EG.5.1 has additional substitutions in its spike protein (namely, Q52H and F456L) compared with XBB.1.5. However, the pathogenicity, transmissibility, and immune evasion properties of clinical isolates of EG.5.1 are largely unknown.

In this study, we used wild-type Syrian hamsters to investigate the replicative ability, pathogenicity, and transmissibility of a clinical EG.5.1 isolate. Our data show that there are no obvious differences in growth ability and pathogenicity between EG.5.1 and XBB.1.5, and both EG.5.1 and XBB.1.5 are attenuated compared to a Delta variant isolate.

We also found that EG.5.1 is transmitted more efficiently between hamsters compared with XBB.1.5. In addition, unlike XBB.1.5, we detected EG.5.1 virus in the lungs of four of six exposed hamsters, suggesting that the virus tropism of EG.5.1 is different from that of XBB.1.5 after airborne transmission.

Finally, we assessed the neutralizing ability of plasma from convalescent individuals and found that the neutralizing activity against EG.5.1 was slightly, but significantly, lower than that against XBB.1.5 or XBB.1.9.2. This suggests that EG.5.1 effectively evades humoral immunity and that the amino acid differences in the S protein of EG.5.1 compared with that of XBB.1.5 or XBB.1.9.2 (i.e., Q52H, R158G, and F456L) alter the antigenicity of EG.5.1.

Our data suggest that the increased transmissibility and altered antigenicity of EG.5.1 may be driving its increasing prevalence over XBB.1.5 in the human population.

## Main text

As of August 2023, over 760 million people have been infected with SARS-CoV-2, with approximately 7 million deaths worldwide (https://covid19.who.int/). After the emergence of the first Omicron subvariant at the end of 2021, numerous Omicron lineages emerged in the human population. In February 2023, the SARS-CoV-2 subvariant XBB.1.5, which is a descendant of the recombinant Omicron lineage XBB, replaced previously dominant Omicron variants globally. Then, in addition to XBB.1.5, several XBB sublineages, such as XBB.1.9.1, XBB.1.16 and XBB.2.3, rapidly spread and circulated throughout the world (https://nextstrain.org/). However, since the end of May 2023, the prevalence of EG.5.1, a descendant of XBB.1.9.2, has been on the rise in north America, Europe and several Asian countries and this variant is currently dominant in China, Hong Kong, and Austria (https://covariants.org/variants/23F.Omicron). This increase in EG.5.1 prevalence suggests a substantial growth advantage over the currently prevailing XBB variants, raising concerns that it may represent the next dominant strain. Given this situation, the World Health Organization (WHO) has added EG.5 to the list of Omicron variants of interest (VOI). EG.5.1 has additional substitutions in its spike protein (namely, Q52H and F456L) compared with XBB.1.5. A recent study evaluated the neutralising activity of plasma from people who experienced breakthrough infection after receiving COVID-19 vaccines by using a pseudotyped virus possessing the EG.5.1 spike protein^1,2^. However, to date, there have been no virological characterization studies *in vitro* or *in vivo* using an authentic EG.5.1 clinical isolate. Accordingly, in this study, we examined the antigenicity, replicative ability, pathogenicity, and transmissibility of EG.5.1 by using a hamster model that is susceptible to SARS-CoV-2.

We amplified the EG.5.1 clinical isolate and confirmed that it contained two additional amino acid changes (i.e., Q52H and F456L), compared to an XBB.1.5 isolate. In addition, our isolate encoded an R158G substitution in the N-terminal domain (NTD) (**Fig. 1**). We first evaluated the pathogenicity of this EG.5.1 isolate in wild-type Syrian hamsters. Intranasal infection with B.1.617.2 (Delta) caused considerable body weight loss (**Fig. 2a**) consistent with our previous observations^3,4^. By contrast, all of the animals infected with the XBB.1.5 or EG.5.1 isolate gained weight for 10 days, similar to the mock-infected animals. We also assessed the growth ability of the viruses in respiratory organs. At 3 and 6 days post-infection (dpi), XBB.1.5 and EG.5.1 replicated in the lungs of the infected animals with no significant differences. However, the viral titres in the nasal turbinates of the EG.5.1-infected hamsters were significantly higher than those in the XBB.1.5-infected hamsters at 6 dpi. In addition, the viral titres in hamsters infected with XBB.1.5 or EG.5.1 were significantly lower than those in hamsters infected with B.1.617.2 at both timepoints (**Fig. 2b**).

**Figure 1.**
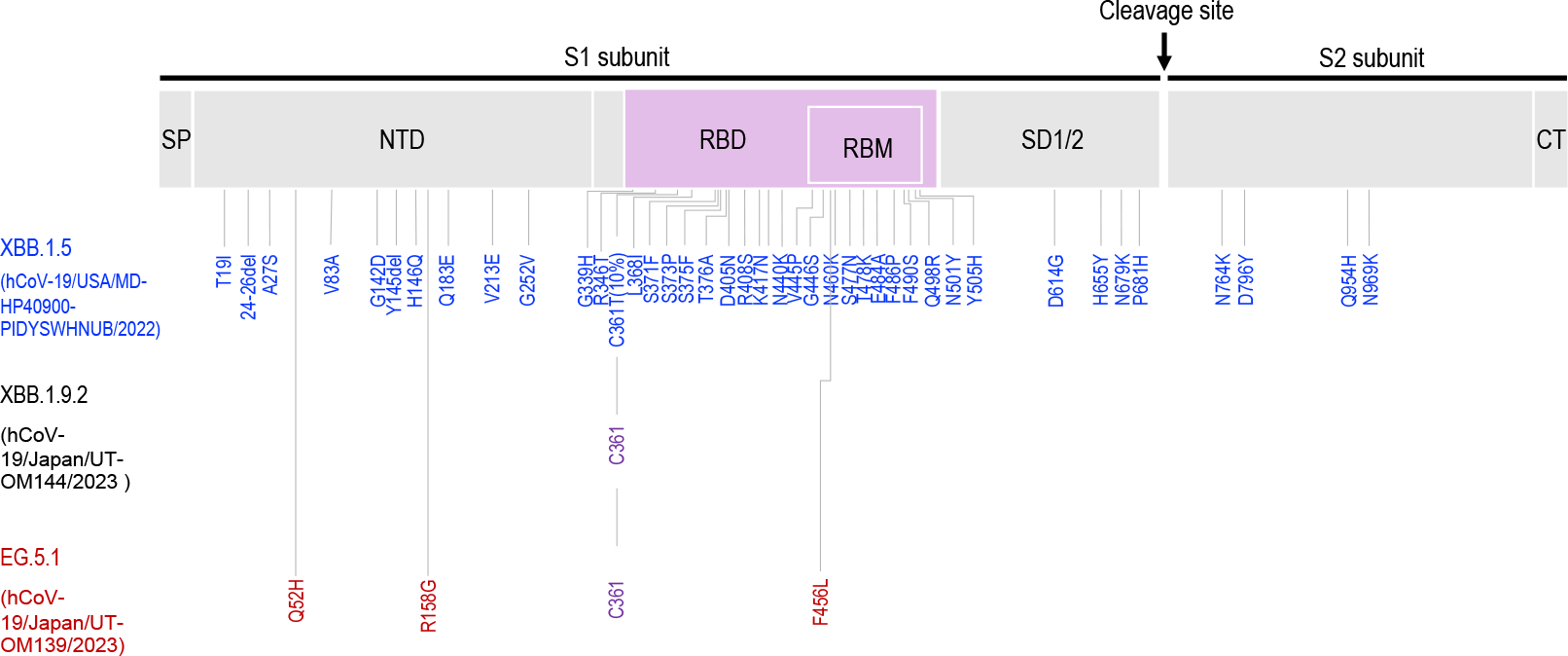
Mutations of Omicron subvariants. Spike (S) protein substitutions in the EG.5.1 and XBB.1.9.2 clinical isolates used in this study. Compared with XBB.1.5 (hCoV-19/USA/MD-HP40900-PIDYSWHNUB/2022), substitutions are shown in red for EG.5.1 (hCoV-19/Japan/UT-OM139/2023). The conserved substitutions between EG.5.1 and XBB.1.9.2 (hCoV-19/Japan/UT-OM144/2023) are shown in purple. The S protein comprises two subunits, S1 and S2. The arrow indicates the S1/S2 proteolytic cleavage site. SP, signal peptide; NTD, N-terminal domain; RBD, receptor-binding domain; RBM, receptor-binding motif; SD1/2, subdomain 1 and 2; CT, cytoplasmic tail.

**Figure 2.**
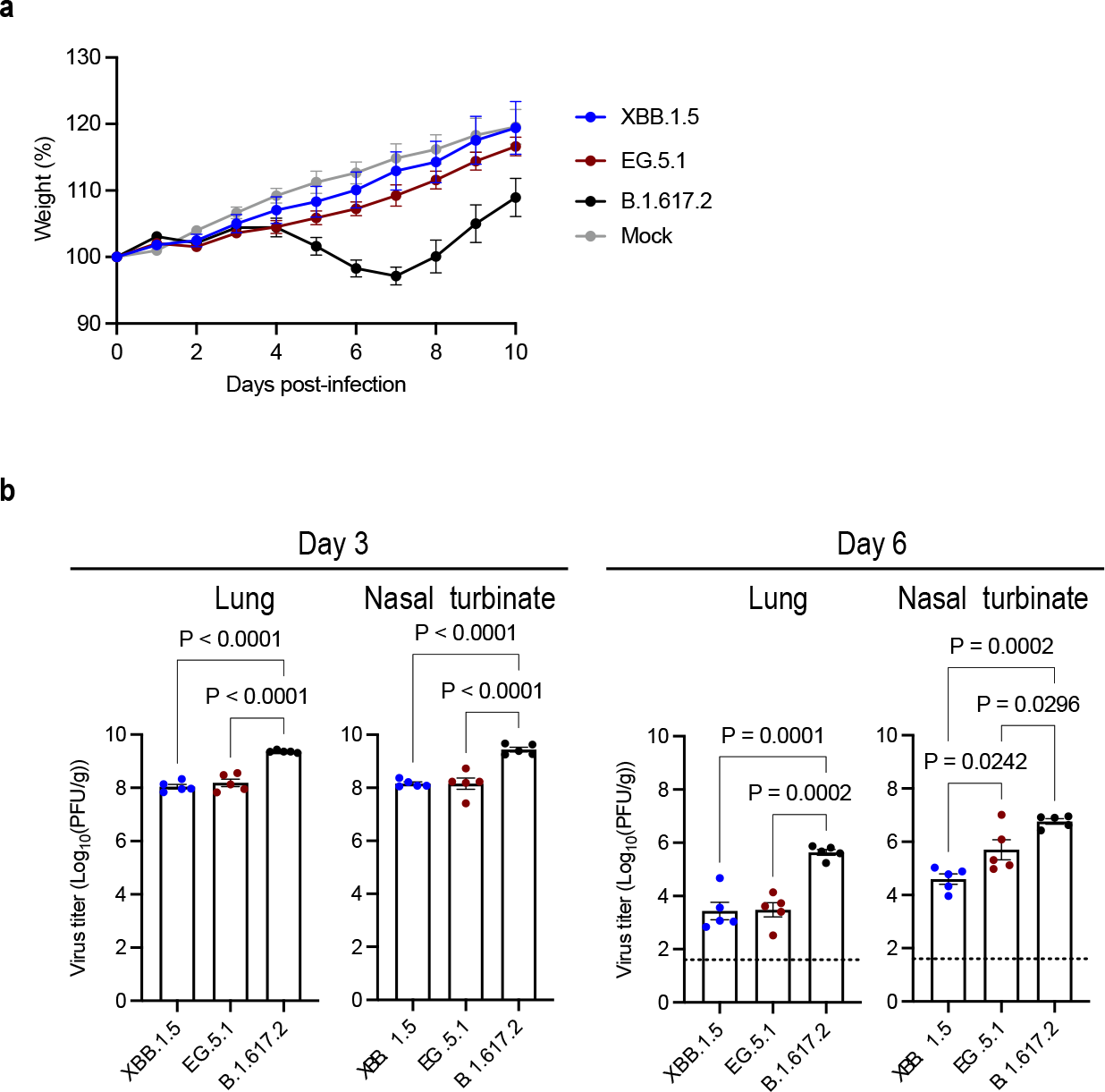
The infectivity and pathogenicity of EG.5.1 in wild-type hamsters. **a**, Wild-type Syrian hamsters were intranasally inoculated with 10^5^ PFU in 30 μL of XBB.1.5 (hCoV-19/USA/MD-HP40900-PIDYSWHNUB/2022) (*n* = 5), EG.5.1 (hCoV-19/Japan/UT-OM139/2023) (*n* =5), B.1.617.2 (hCoV-19/USA/WI-UW-5250/2021) (*n* = 5), or PBS (mock) (*n* = 5). Body weights of virus-infected and mock-infected hamsters were monitored daily for 10 days. Data are presented as the mean percentages of the starting weight (± s.e.m.). **b**, Virus replication in infected Syrian hamsters. Hamsters (*n* =10) were intranasally inoculated with 10^5^ PFU in 30 μL of XBB.1.5 (hCoV-19/USA/MD-HP40900-PIDYSWHNUB/2022), EG.5.1 (hCoV-19/Japan/UT-OM139/2023), or B.1.617.2 (hCoV-19/USA/WI-UW-5250/2021) and euthanized at 3 and 6 dpi for virus titration (*n* = 5/day). Virus titres in the nasal turbinates and lungs were determined by performing plaque assays with Vero E6-TMPRSS2-T2A-ACE2 cells. Vertical bars show the mean ± s.e.m. Points indicate data from individual hamsters. The lower limit of detection is indicated by the horizontal dashed line. Data were analysed by using a one-way ANOVA with Tukey’s multiple comparisons test. *P* values of < 0.05 were considered statistically significant.

We recently reported that XBB.1.5 has better airborne transmissibility than its predecessor, BA.2, which did not transmit at all among hamsters^5^. Given the increasing prevalence of EG.5.1 over XBB.1.5, we evaluated the transmissibility of EG.5.1 in hamsters. Similar to our previous findings, B.1.617.2 transmitted efficiently (100% transmission) ^5^. We previously showed that for XBB.1.5, the virus was detected in the nasal turbinates of five of nine pairs of exposed hamsters (56% transmission), but not in the lungs of any animals^5^. In contrast, unlike XBB.1.5, infectious EG.5.1 virus was detected not only in the nasal turbinates of three of the six EG.5.1-exposed hamsters (#1,2 and 4), but also in the lungs of four of the six exposed hamsters (#1,3,4 and 5) (**Fig. 3**). Of note, in two exposed hamsters (#3 and 4), the infectious virus was detected only in the samples from the lungs. Given the slightly higher transmission efficacy of EG.5.1 (five of six pairs of hamsters: 83%) compared with that of XBB.1.5 (56% transmission) and the detection of virus in the lungs of most of the EG.5.1-exposed animals (four of six exposed animals: 67%) as opposed to no virus detection in the lungs of the XBB.1.5-exposed animals, further evaluation is required to determine what factors contribute to the difference in transmission efficacy and virus tropism between XBB.1.5 and EG.5.1 during airborne transmission.

**Figure 3.**
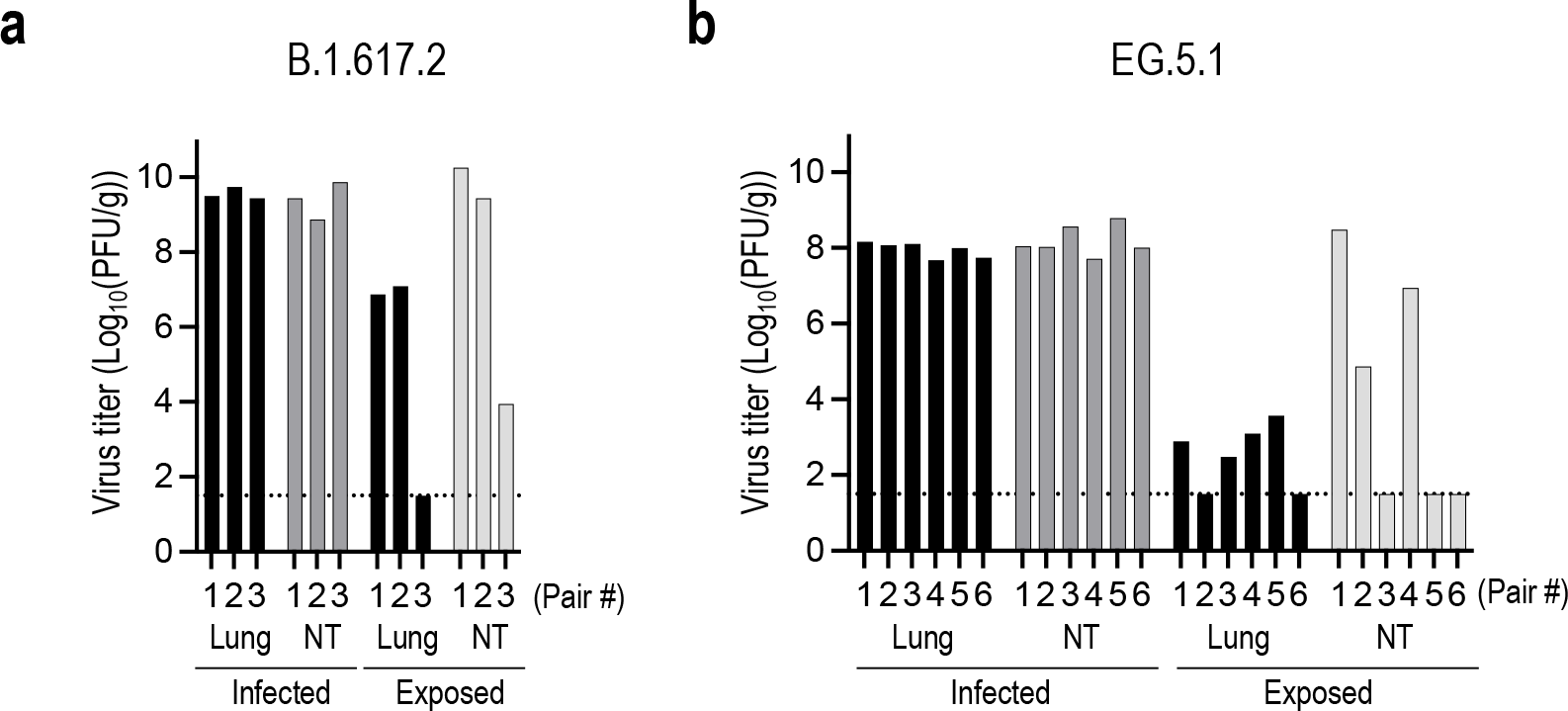
Airborne transmission of B.1.617.2 and EG.5.1 in Syrian hamsters. Virus titres in the lung and nasal turbinate (NT) tissues of infected hamsters and exposed hamsters. Transmission was evaluated for (**a**) B.1.617.2 (3 pairs of hamsters) and (B) EG.5.1 (6 pairs of hamsters). Virus titres are indicated by single bars for each hamster. Virus titres in the nasal turbinates and lungs were determined by performing plaque assays with Vero E6-TMPRSS2-T2A-ACE2 cells. The dotted line indicates the limit of detection.

Lastly, to evaluate the immune evasion of EG.5.1, we tested the neutralising ability against EG.5.1 of plasma from individuals who received mRNA vaccines and experienced breakthrough infections with variants circulating after March 2023 (**Fig. 4, Table 1**). To assess the impact of the amino acid substitutions in EG.5.1, we used XBB.1.5 and XBB.1.9.2 clinical isolates for comparison. Although all tested plasma samples had neutralising activity against EG.5.1, the FRNT_50_ geometric mean titres against XBB.1.5, XBB.1.9.2, and EG.5.1 were 5.4-, 5.4- and 10.2-fold lower than those against the ancestral strain, respectively. Notably, the neutralising activity against EG.5.1 was slightly, but significantly, lower than that against XBB.1.5 or XBB.1.9.2 (**Fig. 4, Table 1**). These results suggest that EG.5.1 effectively evades humoral immunity induced by infection of recently circulating variants including XBB subvariants, and that the amino acid differences in the S protein of EG.5.1 compared with that of XBB.1.5 or XBB.1.9.2 (i.e., Q52H, R158G, and F456L) alter the antigenicity of EG.5.1, leading to its higher immune evasion capability. A recent study has shown that imprinting of humoral immunity reduces the diversity of neutralising antibodies, which suggests that the lower neutralising activity against EG.5.1 after breakthrough infection of recently circulating strains may be influenced by immune imprinting^6^.

**Table 1.** Neutralising antibody titres of human plasma from individuals who were infected with an omicron variant after the end of March after receiving several doses of COVID-19 mRNA vaccines.

| Sample ID | Age | Gender | Plasma collection day post-onset | Infected Omicron sublineage if known | Number of Vaccinations | Remarks | FRNT50: 50% focus reduction neutralization titre |  |  |  |
| --- | --- | --- | --- | --- | --- | --- | --- | --- | --- | --- |
|  |  |  |  |  |  |  | SARS-CoV-2/UT-NC002-1T/Human/2020/Tokyo (A) | hCoV-19/USA/MD-HP40900-PIDYSWHNUB/2022 (Omicron XBB.1.5) | hCoV-19/Japan/UT-OM144/2023 (Omicron XBB.1.9.2) | hCoV-19/Japan/UT-OM139/2023 (Omicron EG.5.1) |
| HP(H)-282 | 44 | M | 12 | Unknown | 5 |  | 623 | 158 | 84 | 37 |
| HP(H)-131 | 62 | M | 40 | Unknown | 5 |  | 2566 | 491 | 498 | 208 |
| HP(H)-129 | 67 | M | 66 | Unknown | 5 |  | 1965 | 551 | 555 | 273 |
| HP(H)-148 | 45 | F | 14 | Unknown | 6 |  | 1456 | 360 | 344 | 176 |
| HP(H)-230 | 36 | M | 14 | Unknown | 5 |  | 526 | 135 | 117 | 52 |
| DACo-001 | 65 | F | 33 | Unknown | 6 |  | 2217 | 984 | 870 | 597 |
| HICo-071 | 75 | M | 104 | Unknown | 4 | malignant tumor | 1138 | 614 | 785 | 365 |
| HICo-073 | 64 | M | 39 | Unknown | 3 | malignant tumor | 2645 | 872 | 865 | 661 |
| HICo-075 | 68 | M | 32 | Unknown | 5 | malignant tumor | 2445 | 613 | 686 | 194 |
| NCCo-1350 | 82 | M | 5 | XBB.1.5 | 3 |  | 409 | <10 | <10 | <10 |
| NCCo-1351 | 76 | M | 4 | XBB.1.5 | 2 |  | <10 | <10 | <10 | <10 |
| NCCo-1355 | 46 | F | 11 | XBB.1.9.1 | 4 |  | 540 | 85 | 88 | 35 |
| NCCo-1356 | 63 | M | 11 | Unknown | 2 |  | 5247 | 648 | 845 | 192 |
| NCCo-1357 | 85 | F | 5 | BN.1.3 | 5 |  | 925 | 22 | 25 | <10 |
| NCCo-1358 | 65 | F | 9 | Unknown | 5 |  | 571 | 27 | 35 | <10 |
| NCCo-1359 | 66 | M | 6 | Unknown | 4 |  | 182 | <10 | <10 | <10 |
| NCCo-1361 | 83 | F | 8 | Unknown | 5 |  | 490 | 182 | 251 | 40 |
| NCCo-1363 | 62 | F | 2 | XBB.1.5 | 2 |  | <10 | <10 | <10 | <10 |
| NCCo-1364 | 28 | F | 4 | XBB.1.9.1 | 4 |  | 82 | <10 | <10 | <10 |
| NCCo-1366 | 71 | M | 10 | Unknown | 4 |  | 5647 | 4629 | 4066 | 2254 |
| NCCo-1371 | 85 | M | 5 | Unknown | 4 |  | 155 | 12 | <10 | <10 |
| NCCo-1377 | 51 | M | 5 | Unknown | 4 |  | 86 | <10 | <10 | <10 |
| NCCo-1378 | 83 | M | 3 | Unknown | 6 |  | 813 | 53 | 45 | 26 |

**Figure 4.**
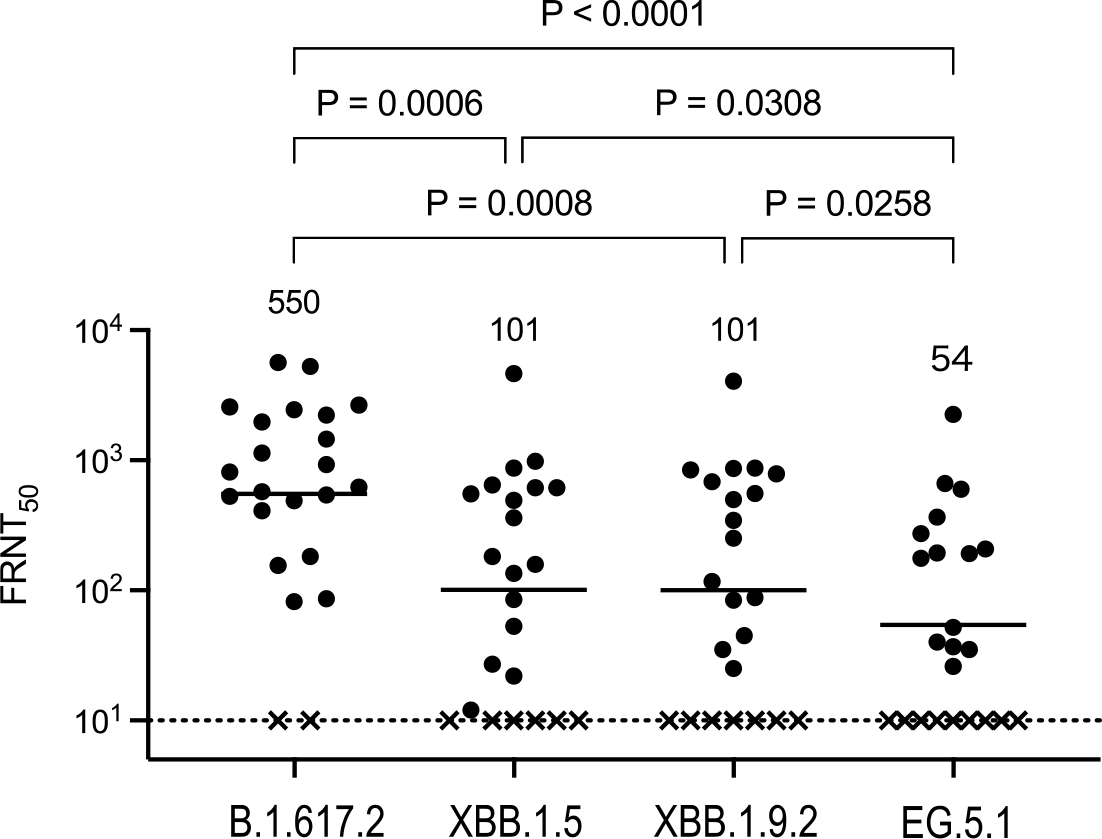
*In vitro* neutralising activity of plasma against SARS-CoV-2 Omicron variants. Neutralising titres of plasma samples obtained from individuals who were infected with an omicron variant after the end of March after receiving several does of mRNA vaccines (n=23). Detailed information about the participants is provided in Tables S1. FRNT_50_ values were determined in Vero E6-TMPRSS2-T2A-ACE2 cells. Each dot represents data from one individual. The lower limit of detection (value=10) is indicated by the horizontal dashed line. Samples under the detection limit (<10-fold dilution) were assigned an FRNT_50_ of 10 and are represented by X. Geometric mean titres are shown. Data were analysed by using a Friedman test followed by Dunn’s test. *P* values of < 0.05 were considered statistically significant

Overall, although EG.5.1 has similar replicative ability in naive wild-type hamsters as XBB.1.5, the transmissibility of EG.5.1 is slightly higher than that of XBB.1.5. In addition, the virus tropism of EG.5.1 is different from that of XBB.1.5 after airborne transmission, since transmitted EG.5.1 virus was detected in not only the nasal turbinates but also the lungs of hamsters. The immune-evading properties of EG.5.1 are slightly but significantly enhanced relative to those of XBB.1.5 and its predecessor XBB.1.9.2. Therefore, the increased transmissibility and altered antigenicity of EG.5.1 may be driving its increasing prevalence over XBB.1.5 in the human population.

## Materials and Methods

### Cells

Vero E6-TMPRSS2-T2A-ACE2 cells were cultured in Dulbecco’s modified Eagle’s medium (DMEM) containing 10% Fetal Calf Serum (FCS), 100 U/mL penicillin–streptomycin, and 10 µg/mL puromycin. VeroE6/TMPRSS2 (JCRB 1819) cells were propagated in the presence of 1 mg/ml geneticin (G418; Invivogen) and 5 μg/ml plasmocin prophylactic (Invivogen) in DMEM containing 10% FCS. Vero E6-TMPRSS2-T2A-ACE2 and VeroE6/TMPRSS2 cells were maintained at 37 °C with 5% CO_2_. The cells were regularly tested for mycoplasma contamination by using PCR, and confirmed to be mycoplasma-free.

### Viruses

The SARS-CoV-2 viruses hCoV-19/Japan/UT-OM139/2023 (Omicron EG.5.1; isolated using Vero E6-TMPRSS2-T2A-ACE2 cells), hCoV-19/Japan/UT-OM144/2023 (Omicron XBB.1.9.2; isolated using Vero E6-TMPRSS2-T2A-ACE2 cells), hCoV-19/USA/MD-HP40900-PIDYSWHNUB/2022 (Omicron XBB.1.5)^7^, hCoV-19/USA/WI-UW-5250/2021 (B.1.617.2, Delta) and SARS-CoV-2/UT-NC002-1T/Human/2020/Tokyo (ancestral strain) were propagated in VeroE6/TMPRSS2 cells.

All experiments with SARS-CoV-2 were performed in enhanced biosafety level 3 containment laboratories at the University of Tokyo and the National Institute of Infectious Diseases, Japan, which are approved for such use by the Ministry of Agriculture, Forestry, and Fisheries, Japan.

### Clinical specimens

After informed consent was obtained, plasma specimens were collected from COVID-19 convalescent individuals (breakthrough infection). The research protocol was approved by the Research Ethics Review Committee of the Institute of Medical Science of the University of Tokyo (approval numbers: 2019–71–0201 and 2020-74-0226).

### Animal experiments and approvals

Animal studies were carried out in accordance with the recommendations in the Guide for the Care and Use of Laboratory Animals of the National Institutes of Health. The protocols were approved by the Animal Experiment Committee of the Institute of Medical Science, the University of Tokyo (approval number PA19-75). Virus inoculations were performed under isoflurane, and all efforts were made to minimize animal suffering.

### Focus reduction neutralisation test (FRNT)

Neutralisation activities of human plasma were determined by using a focus reduction neutralisation test as previously described^8^. The samples were first incubated at 56 °C for 1 h. Then, the treated plasma samples were serially diluted five-fold with DMEM containing 2% FCS in 96-well plates and mixed with 100–400 FFU of virus/well, followed by incubation at 37 °C for 1 h. The plasma-virus mixture was inoculated onto Vero E6-TMPRSS2-T2A-ACE2 cells in 96-well plates in duplicate. After a 1-h incubation at 37 °C, 100 μl of 1.5% Methyl Cellulose 400 (FUJIFILM Wako Pure Chemical Corporation) in culture medium was then added to each well. The cells were incubated for 14–18 h at 37 °C and then fixed with formalin.

After the formalin was removed, the cells were immunostained with a mouse monoclonal antibody against SARS-CoV-2 nucleoprotein [N45 (TAUNS Laboratories, Inc., Japan)], followed by a horseradish peroxidase-labeled goat anti-mouse immunoglobulin (Jackson ImmunoResearch Laboratories Inc.). The infected cells were stained with TrueBlue Substrate (SeraCare Life Sciences) and then washed with distilled water. After cell drying, the focus numbers were quantified by using an ImmunoSpot S6 Analyzer, ImmunoCapture software, and BioSpot software (Cellular Technology). The results are expressed as the 50% focus reduction neutralisation titre (FRNT_50_). The FRNT_50_ values were calculated by using GraphPad Prism (GraphPad Software). Samples under the detection limit (<10-fold dilution) were assigned an FRNT_50_ value of 10.

### Experimental infection of Syrian hamsters

For virological and pathological examinations, under isoflurane anesthesia, five six-week-old male wild-type Syrian hamsters (Japan SLC Inc., Shizuoka, Japan) per group were intranasally inoculated with 10^5^ PFU (in 30 μL) of EG.5.1 (hCoV-19/Japan/UT-OM139/2023), XBB.1.5 (hCoV-19/USA/MD-HP40900-PIDYSWHNUB/2022), or B.1.617.2 (hCoV-19/USA/WI-UW-5250/2021). Baseline body weights were measured before infection. Body weight was monitored daily for 10 days. To evaluate virus growth capability in hamsters, ten hamsters per group were intranasally infected with 10^5^ PFU (in 30 μL) of EG.5.1 hCoV-19/Japan/UT-OM139/2023), XBB.1.5 (hCoV-19/USA/MD-HP40900-PIDYSWHNUB/2022), or B.1.617.2 (hCoV-19/USA/WI-UW-5250/2021); 3 and 6 dpi, five animals were euthanized and nasal turbinates and lungs were collected. The virus titres in the nasal turbinates and lungs were determined by use of plaque assays on Vero E6-TMPRSS2-T2A-ACE2 cells.

For the airborne transmission study between hamsters, six-week-old male hamsters were intranasally inoculated with 10^5^ PFU (in 30 μL) of EG.5.1 (hCoV-19/Japan/UT-OM139/2023) or B.1.617.2 (hCoV-19/USA/WI-UW-5250/2021) while under isoflurane anesthesia. Infected donor hamsters were housed in wire cages inside an isolator unit. Twenty-four hours later, naïve hamsters were placed on the other side of the cage. A double-layered wire mesh separated the hamsters by 5 cm to prevent direct contact. The infected, donor hamsters were positioned in the front of the isolator unit, which provided unidirectional airflow. Tissue samples were collected 3 days after infection for the donor hamsters or 3 days after initial contact for the exposed hamsters. The virus titres in the nasal turbinates and lungs were determined by use of plaque assays on Vero E6-TMPRSS2-T2A-ACE2 cells.

### Whole genome sequencing

Viral RNA was extracted by using a QIAamp Viral RNA Mini Kit (QIAGEN). The whole genome of SARS-CoV-2 was amplified by using a modified ARTIC network protocol in which some primers were replaced or added. Briefly, viral cDNA was synthesized from the extracted RNA by using a LunarScript RT SuperMix Kit (New England BioLabs). The DNA was amplified by performing a multiplexed PCR in two pools using the ARTIC-N6 primers and the Q5 High-Fidelity DNA polymerase or Q5 Hot Start DNA polymerase (New England BioLabs). The DNA libraries for Illumina NGS were prepared from pooled amplicons by using a QIAseq FX DNA Library Kit (QIAGEN) and were then analysed by using iSeq 100 System (Illumina). To determine the virus sequences, the reads were assembled by CLC Genomics Workbench (version 23, Qiagen) with the Wuhan/Hu-1/2019 sequence (GenBank accession no. MN908947) as a reference. The sequences of Omicron EG.5.1 (hCoV-19/Japan/UT-OM139/2023) and Omicron XBB.1.9.2 (hCoV-19/Japan/UT-OM144/2023) were deposited in the Global Initiative on Sharing All Influenza Data (GISAID) database with accession IDs EPI_ISL_18134846 and EPI_ISL_18134847, respectively.

### Statistical analysis

GraphPad Prism was used to analyse all of the data. Statistical analysis included a one-way ANOVA with Tukey’s multiple comparisons test, and the Friedman test followed by Dunn’s test. Differences among groups were considered significant for *P* < 0.05.

## Acknowledgements and funding

We thank Susan Watson for scientific editing. We also thank the Medical Association of Kashiwa for collecting clinical samples. In addition, we thank Kyoko Yokota, Tomoka Nagashima, Rie Onoue, and Mao Suzuki for technical assistance. Vero E6-TMPRSS2-T2A-ACE2 cells were provided by Dr. Barney Graham, NIAID Vaccine Research Center.

This study was supported by grants from the Center for Research on Influenza Pathogenesis and Transmission (75N93021C00014), by the National Institute of Allergy and Infectious Diseases, and a research programme on emerging and re-emerging infectious diseases (JP21fk0108552), the Japan Program for Infectious Diseases Research and Infrastructure (JP23wm0125002), and the Japan Initiative for World-leading Vaccine Research and Development Centers (JP233fa627001) from the Japan Agency for Medical Research and Development. The funders had no role in the study design, data collection, data analysis, interpretation, or writing of the paper.

## Author Contributions

R.U.: conceptualization, data curation, formal analysis, validation, visualization, and writing of the first draft. M. Kiso: data curation, formal analysis, and methodology. K.I-H.: data curation, resources and validation. S.Y.: conceptualization, data curation, formal analysis, and methodology. M. Ito and S.C.: data curation, Y.S-T., M. Imai, M. Koga, N.F., N.Okumura., M.H., N.I, H.K., H.N., N.Ohmagari., and H.Y.: resources. Y.Kashima., and Y.S.: virus isolation and data curation. Y. Kawaoka: conceptualization, supervision, writing (review and editing), and funding acquisition. R.U., M. Kiso, K.I-H., and S. Y. contributed equally.

